# Epigenetic findings in periodontitis in UK twins: a cross sectional study

**DOI:** 10.1101/381327

**Authors:** Yuko Kurushima, Pei-Chien Tsai, Juan Castillo Fernandez, Alexessander Couto Alves, Julia Sarah El-Sayed Moustafa, Caroline Le Roy, Mark Ide, Francis J Hughes, Kerrin Small, Claire J Steves, Jordana T Bell

## Abstract

**Background:** Genetic and environmental risk factors contribute to periodontal disease, but the underlying susceptibility pathways are not fully understood. Epigenetic mechanisms are malleable regulators of gene function that can change in response to genetic and environmental stimuli, thereby providing a potential mechanism for mediating risk effects in periodontitis. The aim of this study is to identify epigenetic changes across tissues that are associated with periodontal disease.

**Methods:** Self-reported gingival bleeding and history of gum disease, or tooth mobility, were used as indicators of periodontal disease. DNA methylation profiles were generated using the Infinium HumanMethylation450 BeadChip in whole blood, buccal, and adipose tissue samples from predominantly older female twins (mean age 58) from the TwinsUK cohort. Epigenome-wide association scans (EWAS) of gingival bleeding and tooth mobility were conducted in whole blood in 528 and 492 twins, respectively. Subsequently, targeted candidate gene analysis at 28 genomic regions was carried out testing for phenotype-methylation associations in 41 (tooth mobility) and 43 (gingival bleeding) buccal, and 501 (tooth mobility) and 556 (gingival bleeding) adipose DNA samples.

**Results:** Epigenome-wide analyses in blood identified one CpG-site (cg21245277 in *ZNF804A*) associated with gingival bleeding (FDR=0.03, nominal p-value=7.17e-8), and 58 sites associated with tooth mobility (FDR<0.05) with the top signals in *IQCE* and *XKR6*. Epigenetic variation at 28 candidate regions (256 CpG-sites) for chronic periodontitis showed a strong enrichment for association with periodontal traits, and signals in eight genes (*VDR, IL6ST, TMCO6, IL1RN, CD44, IL1B, WHAMM*, and *CXCL1*) were significant in both traits. The methylation-phenotype association signals validated in buccal samples, and a subset (25%) also validated in adipose tissue.

**Conclusions:** Epigenome-wide analyses in adult female twins identified specific DNA methylation changes linked to self-reported periodontal disease. Future work will explore the environmental basis and functional impact of these results to infer potential for strategic personalized treatments and prevention of chronic periodontitis.

## BACKGROUND

Periodontal disease is a chronic inflammatory disease that poses a significant burden among older populations. The disease causes inflammation and bleeding of the gingival tissues as a result of the accumulation of microbial dental plaque, and in its destructive form, periodontitis, results in progressive destruction of the supporting structures of teeth leading to looseness of the teeth and ultimately tooth loss. The aetiology of the disease involves interplay between both intrinsic and extrinsic factors such as genetic susceptibility, diet, oral hygiene, and inflammatory response to exposure to complex pathogens. An increasing number of genome-wise association studies [1] [2] and candidate-gene reports [3] [4] [5] have explored genetic susceptibility to periodontal disease. Genes that harbor genetic variants robustly associated with the development of periodontal disease are related to inflammatory response and alveolar bone resorption; however, their mechanistic pathways are not clearly understood.

Epigenetic modifications, including DNA methylation, are key regulators of gene function. Epigenetic marks can change in response to genetic and environmental stimuli, thereby providing a mechanism for the interaction of genetic and environmental effects in the context of disease susceptibility. A number of epigenetic changes have been identified to contribute towards and act as biomarkers of human diseases, including cancers, inflammatory and neurological disorders, and mental health traits [6] [7]. However, only a few studies so far have explored epigenetics in periodontal disease, although periodontal disease is characterized by an inflammatory response. The majority of epigenetic research into periodontal disease to date has focused on candidate genes in case-control samples [8] [9] [10]. Previous studies predominantly explored the hypothesis that epigenetic regulation mediates gingival inflammation in epithelial tissue by changing the function of cytokine genes. This includes specifically the up-regulation of proinflammatory cytokines and other signaling molecules, and down-regulation of anti-inflammatory cytokines to activate a full inflammatory response.

Although both genetic and environmental risk factors contribute to periodontal disease susceptibility, the underlying molecular pathways of these risk effects are not fully understood. Epigenetic studies may uncover mechanisms mediating genetic and environmental risk effects on periodontitis, which will help towards developing strategic personalized treatments and prevention.

The aim of the current study was to investigate regulatory genomic variation associated with chronic periodontal disease using a two-fold approach. The first stage explored the association of periodontal traits with epigenome-wide variation in over 500 female adult twins. Thereafter, a candidate gene approach was undertaken, whereby we explored multi-omics signatures at candidate genomic regions previously implicated in periodontitis [1] [2] [11] [12] [13]. The association between periodontal status and DNA methylation levels was assessed in whole blood, as the most widely used sample type, in adipose tissue as an independent sample type, and in buccal tissue, as a representative biological sample from the oral cavity.

## MATERIALS AND METHODS

### Study population

Study participants were older female twins (mean age = 58 years old; age range: 19-82 years old; 83% are MZ and 10% are DZ) who were enlisted in the TwinsUK registry [14]. The TwinsUK registry is one of the largest adult twin cohorts encompassing a wide range of phenotypes, and biological samples and data. Informed consent was obtained from all participants before sample and data collection. Zygosity was determined by standard questionnaire [15] and confirmed by DNA short-tandem repeat fingerprinting. Gingival bleeding status was obtained from 528 female individuals (mean age 57.9), and tooth mobility data was obtained from 492 female individuals (mean age 58.0), for whom blood DNA methylation profiles were also available. The date of the clinical visit for periodontal data collection was within 5 years of the blood DNA extraction date. Buccal tissue DNA methylation samples included 43 female individuals (mean age 57, age range 34-73; subjects including 20 MZ individuals, 16 DZ individuals, and 7 singletons) with gingival bleeding phenotype and 41 females (mean age 57, range 34-73, 20 MZ individuals, 12 DZ individuals, 9 singletons) with tooth mobility phenotype. Lastly, the adipose tissue DNA methylation sample included 556 female individuals (mean age 59, range 39-85, 39% MZ) with gingival bleeding phenotype and 501 female individuals (mean age 58, range 39-85, 39% MZ, 61%DZ) with tooth mobility phenotype.

### Periodontal phenotypes

Periodontal phenotype data on tooth mobility and gingival bleeding were collected by self-reported questionnaires during clinical visits. A single question was used to determine each of gingival bleeding (question of “Have you ever had the condition of gum bleeding”) and tooth mobility (question of “Have you ever had the condition of gum decay or loose teeth”). Participants were asked to answer either “yes” or “no” to these questions. Therefore both phenotypes were binary variables. In the blood DNA methylation dataset, 269 individuals out of 528 (51%) were gingival bleeding positive and 121 out of 492 (25%) were tooth mobility positive. In the buccal methylation dataset, 25 individuals out of 43 were gingival bleeding positive and 12 individuals out of 41 were tooth mobility positive. In the adipose DNA methylation dataset, 265 individuals out of 556 were gingival bleeding positive and 102 individuals out of 501 were tooth mobility positive.

### Epigenome-wide profiling of DNA methylation

The Infinium HumanMethylation450 BeadChip Kit (Illumina Inc., San Diego, CA, USA) was used to obtain the ratio of array intensity signal from the methylated beads over the sum of methylated and unmethylated bead signals plus 100. This array contains a combination of two types of probes, Infinium I (28%) and Infinium II (72%) probes, which use different detection methods. Infinium I probes include two bead types, a methylated and an unmethylated, while Infinium II probes are measured in one bead type that requires extension of color labeled dideoxynucleotides to determine the methylation status. To correct for the technical bias causes by the probe design in this platform, the beta mixture quantile normalization (BMIQ) method and background correction was applied for each sample [16]. Subjects with more than 1% of missing values were considered as outliers and were removed from the analysis. Probes with more than 1% of missing values among all samples with detection P-value > 0.05, and probes that mapped incorrectly or that mapped to multiple locations in the genome (with up to 2 mismatches), were removed. We also used the 65 designed single nucleotide polymorphism (SNPs) to check for sample swaps.

In total, after all exclusions we analysed 452874 probes in blood, 436814 probes in buccal tissue, and 467928 probes in adipose samples. The methylation levels at each probe were normalized to standard normal distribution (N (0,1)) before downstream epigenome-wide association analysis. Epigenome-wide association scans aimed to identify specific sites across the genome where epigenetic levels significantly differed between periodontal disease positive and negative group samples, that is, identifying differentially methylated positions (DMPs) for periodontal disease.

### RNA-sequencing gene expression quantification in whole blood

RNA-sequencing (RNASeq) data from peripheral whole blood samples were available for 384 subjects, and were generated, quantified and normalized as previously described [17] [18]. Briefly, RNASeq reads were aligned to the UCSC (University of California Santa Cruz) Genome Browser GRCh37 reference genome with the Burrows-Wheeler Aligner [19], and genes annotated using GENCODE v.10. Samples with fewer than 10 million reads mapping to known exons were excluded. All read count quantifications were carried out at the exon level, and were corrected for variation in sequencing depth between samples by normalizing the number of reads to the median number of well-mapped reads. Exons expressed in fewer than 90% of individuals were excluded. Rank-based inverse normal transformation was then applied to the normalized read counts prior to downstream analysis. The number of individuals included in the gene expression analysis with available gingival bleeding and with tooth mobility phenotypes was 342 and 335, respectively.

### Metabolomics data

Fasting whole blood samples were extracted and stored at −80°C until they were processed by Metabolon, Inc. (Durham, NC, USA) using a non-targeted mass spectrometry platform, as previously described [20]. After profiling, normalization was performed to correct for variation resulting from instrument inter-day tuning differences. Briefly, each compound was normalized so that the median of the run-day was equal to one (1.00) and each data point was scaled proportionally. The analysis only focused on 271 known metabolites, which included amino acids, carbohydrates, vitamins, lipids, nucleotides, small peptides and xenobiotics. This metabolomics data was available in 472 (tooth mobility) and 506 (gingival bleeding) individuals for whom phenotype and whole blood DNA methylation data were also profiled. The overlap in datasets across the two periodontal traits, DNA methylation, and blood gene expression and metabolomics samples is shown in Table 1.

**Table 1.** Study samples used in blood omics data and methylation profiles in buccal and adipose tissues for two periodontal traits

|  | Gingival bleeding <sup>a)b)</sup> | Tooth mobility | Overlap between two periodontal traits |
| --- | --- | --- | --- |
| Blood |  |  |  |
| DNA methylation | 528 | 492 | 474 |
| RNAseq | 342 (120) | 335 (112) | 333 |
| Metabolomics | 506 (506) | 472 (472) | 454 |
| Buccal DNA methylation | 43 | 41 | 40 |
| Adipose DNA methylation | 556 | 501 | 501 |

a) Units represent the number of individuals.
b) Numbers in brackets represent the number of individuals who also have DNA methylation data

### Candidate gene selection

We focused on 37 candidate genes for chronic periodontitis (Table 2). Some of these genes were selected as they were previously identified through genome-wide association studies (GWAS) of periodontitis, while other regions were included because of their biological function and previous research exploring their contribution to periodontal disease (see Table 2). Each candidate genomic region was defined by the following criteria: for GWAS SNPs the region was set with a range of 10kb around the lead SNP, and for biological candidate genes we included the gene body and 5kb either side of the gene boundary. In total, 28 of the 37 candidate genes were represented by 256 CpG sites on the Infinium HumanMethylation450 array.

**Table 2.**
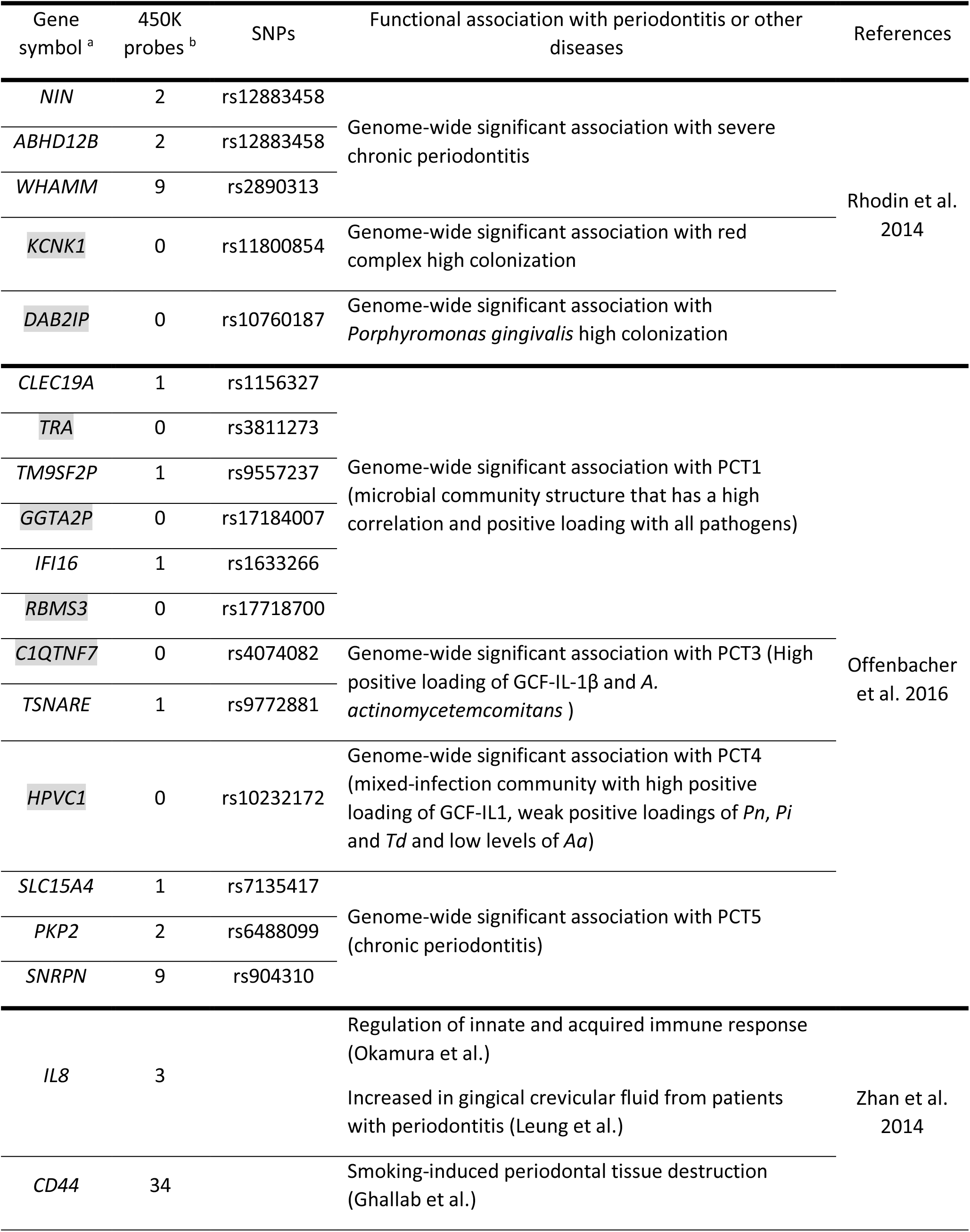
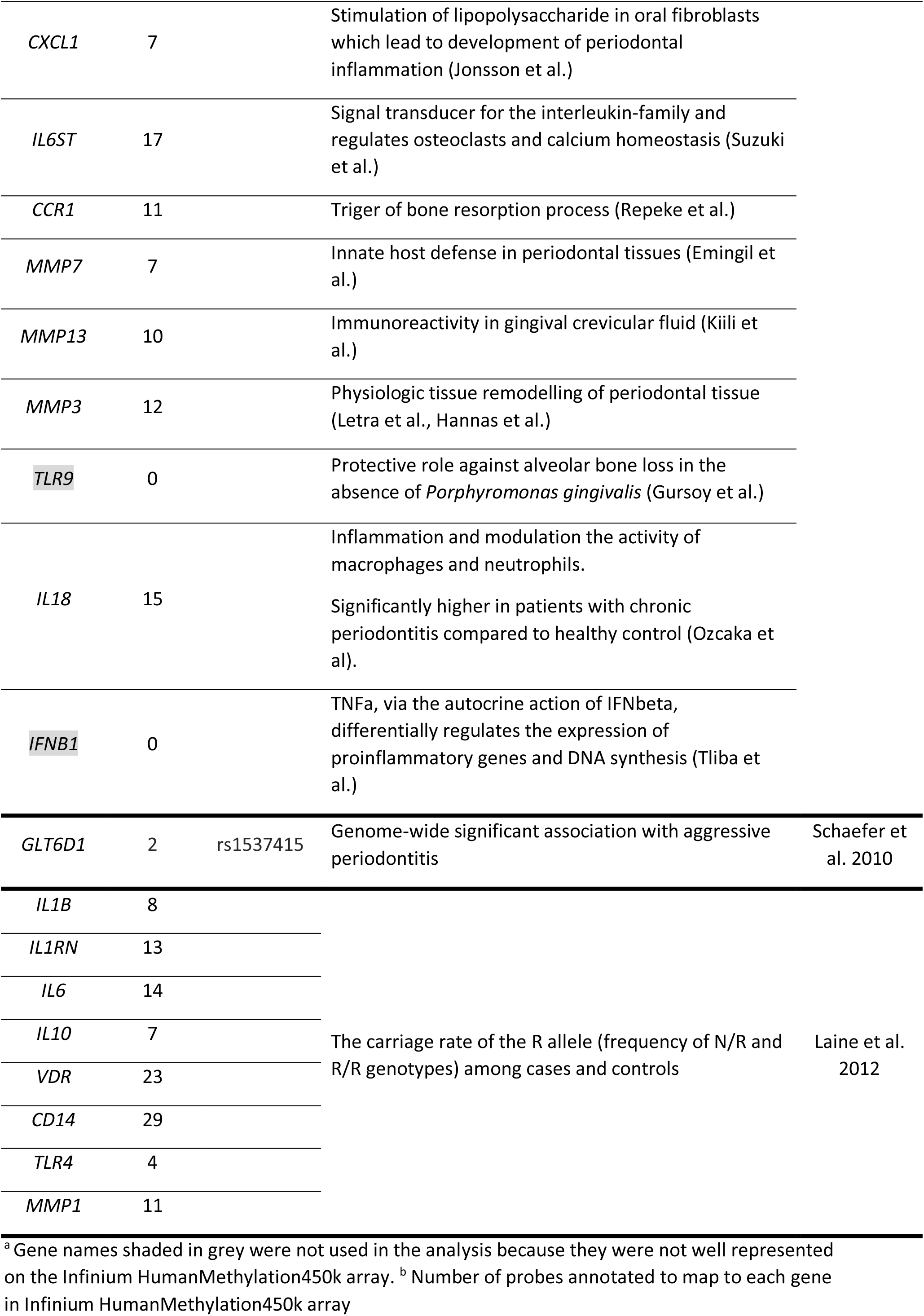
Chronic periodontitis GWAS loci and candidate genes from previous studies.

Further to the epigenome-wide analyses in whole blood, we assessed the association between buccal tissue DNA methylation at the candidate CpG-sites and periodontal traits, and adipose tissue DNA methylation. Buccal DNA methylation analyses were carried out in 41 (tooth mobility) and 43 (gingival bleeding) individuals at a subset of 179 CpG-sites (from the 256) that passed quality control assessment in the buccal dataset.

### Statistical analysis

We performed a linear mixed effects regression (LMER) model to assess the association between DNA methylation levels at each probe and periodontal status of the individual. Normalised DNA methylation levels were the response variable and each periodontal phenotype was a fixed effects predictor, along with a number of covariates. Analyses of blood DNA methylation profiles included fixed effects (age, beadchip, position, BMI, smoking, blood cell composition (monocytes, natural killer cells, granulocytes, CD8 T cells, and CD4 T cells), DNA extraction year, difference between the time point of dental phenotype collection and DNA extraction) and random-effect (family structure and zygosity) variables. The association between DNA methylation levels and periodontal traits were carried out using lmer within R 3.4.3 in the package “Ime4”. In buccal samples, the association between periodontal status and methylation levels was also assessed using LMER adjusting for fixed effects (age, zygosity, family structure and beadchip) and random effects as in blood. In adipose samples fixed effects included age, zygosity, family structure, beadchip, BMI, bisulfite conversion level and bisulfite efficiency, and random effects were as described in blood. Multiple testing correction was carried out using a false discovery rate (FDR) at 5% [21]. Similarly, we tested if exon expression levels were differentially associated with periodontal traits using a linear mixed effects regression model. The analysis was conducted for each exon adjusting for fixed effects (age, age squared, smoking and GC mean, insert size mode) and random effects (family structure, zygosity, date and primer index).

Heritability estimation was conducted using a classical twin study model. This model allows us to estimate the extent of genetic, shared environment and unique environmental contributions to the phenotypic variance [22]. Shared environmental factors represent the familial exposures which twins often share in childhood, whereas unique environmental factors are individual exposures. We calculated heritability in R with the “mets” package.

## RESULTS

### *ZNF804A* methylation associates with gum bleeding

Epigenome-wide analysis of gingival bleeding was carried out in the blood methylome across 452874 probes. We identified one CpG site significantly associated with gingival bleeding (cg21245277; beta = −0.33, p-value = 7.17e-8, FDR = 0.03; Table 3). This CpG site is located within the 5’UTR of a protein coding gene, *ZNF804A* (Figure 1A). Polymorphisms of this gene confer increased susceptibility to mental health disorders such as schizophrenia and bipolar disorders, and contribute to heroin addiction [23]. CpG-site cg21245277 was significantly hypo-methylated in participants who experienced gingival bleeding (Figure 1B).

**Figure 1.**
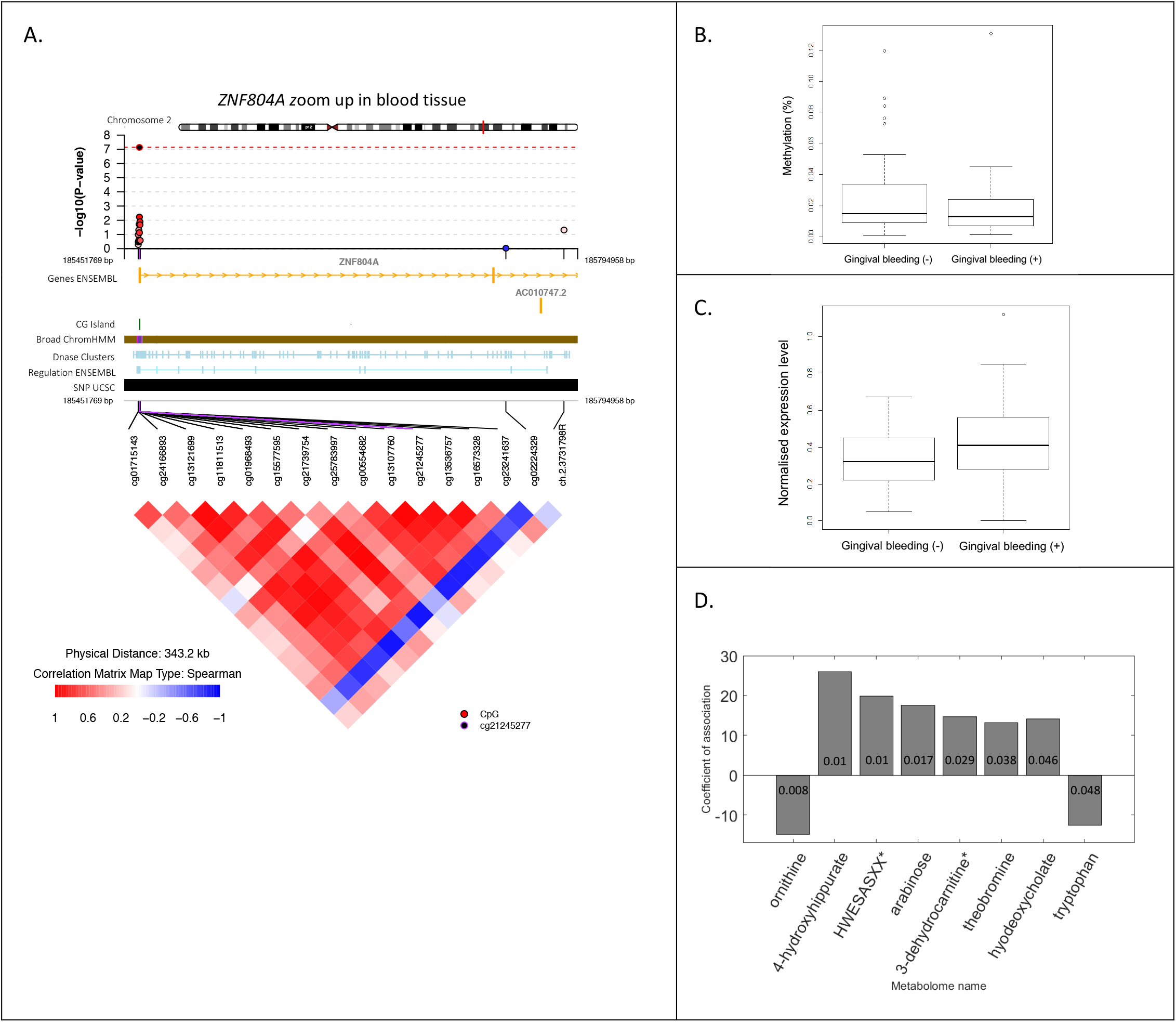
*ZNF804A* differentially methylated signal for gingival bleeding in whole blood. **(A)** Visualisation of the peak blood epigenome-wide signal for gingival bleeding (top), along with genomic annotation of the region (middle) and co-methylation patterns (bottom panel). **(B)** The association between *ZNF804A* blood tissue methylation at cg21245277 and gingival bleeding. The y axis shows uncorrected methylation signals. **(C)** The association between the normalised expression levels at exon ENSG00000170396.6_185463093_185463797 in *ZNF804A* and gingival bleeding (p-value = 0.018). **(D)** The association between *ZNF804A* blood methylation levels at cg21245277 and eight metabolites, which were nominally significant in a multiple correction test across 271 blood metabolites. The y-axis represents the coefficient of the model, and the p-value of the association is shown in each bar.

**Table 3.** The most-associated epigenome-wide significant signals for both periodontal traits

|  | bp | Annotation | Estimate <sup>a)</sup> | P-value | FDR | Gene <sup>b)</sup> | Reported traits |
| --- | --- | --- | --- | --- | --- | --- | --- |
| Gingival<br>bleeding | CHR2:<br>185463218 | 1<sup>st</sup> Exon;<br>5'UTR | -0.33 | 7.17e-8 | 0.03 | ZNF804A | Bipolar Disorder |
|  |  |  |  |  |  | (Zinc Finger<br>Protein 8) | Schizophrenia<br>Memory loss |
| Tooth<br>mobility | CHR7:<br>2646478 | Body | 0.38 | 6.85e-8 | 2.5e-4 | IQCE | Longevity |
|  |  |  |  |  |  | (IQ Motif<br>Containing E) | Blood<br>metabolite level |
|  | CHR8:<br>11058145 | Body | -0.49 | 1.53e-8 | 2.8e-3 | XKR6 | Asthma |
|  |  |  |  |  |  | (XK Related 6) | Allergic rhinitis |

a) Estimate represents the coefficient generated from the epigenetic linear regression model
b) Gene name derived from UCSC

To assess if genetic or environmental factors influence blood DNA methylation levels at this site, we carried out heritability analysis using the classical twin model. We established a moderate impact of genetic variation on DNA methylation variability at this CpG site (heritability of 23%), while the majority (77%) of methylation variance was attributed to environmental factors unique to each individual.

To explore the potential functional role of the methylation signal in *ZNF804A*, we carried out additional ‘omic analyses. First, RNA-sequencing gene expression data in 342 twins identified two exons in *ZNF804A*, which are expressed in blood. One exon showed a nominally significant association with the presence of gingival bleeding (beta = 2.17e-6, *p*-value = 0.018; Figure 1C) in a model adjusting for age, smoking status, family structure and technical covariates. Subjects who experienced gingival bleeding had lower levels of DNA methylation in the 5’UTR of this gene, and higher levels of exon expression compared to controls, suggesting negative epigenetic regulatory effects of cg21245277 on exon expression (correlation = −0.13).

To further explore the functional relevance of the most-associated gingival bleeding DNA methylation signal in *ZNF804A*, we correlated the methylation levels at the most-associated CpG-site in this gene (cg21245277) with 271 known fasting blood metabolites profiled by mass spectrometry [20]. Eight metabolites were found to be nominally associated with *ZNF804A* cg21245277 blood DNA methylation levels (Figure 1D). The strongest signal was observed for ornithine (*p*-value = 0.008), which is also significantly associated with tooth mobility (*p*-value = 0.012). The remaining metabolites also included 3-dehydrocarnitine, ranked fifth (*p*-value = 0.029), which is consumed by gram negative and positive bacteria in either aerobic or anaerobic environmental insults or as a sole carbon, nitrogen and energy source [24].

### Epigenome-wide analyses of self-reported periodontitis

Epigenome-wide analyses of self-reported periodontitis (tooth mobility) identified 58 epigenome-wide significant CpG sites (Figure 2A), although the results showed evidence for moderate genomic inflation (lambda = 1.27). The two most-associated CpG sites (Table 3) were located in the gene body of the *IQCE* gene (cg08157914; beta = 0.38, *p*-value = 6.85e-8, FDR <0.001,) and in the gene body of the *XKR6* gene (cg11051055; beta = −0.49, *p*-value = 1.53e-8, FDR = 0.003).

**Figure 2.**
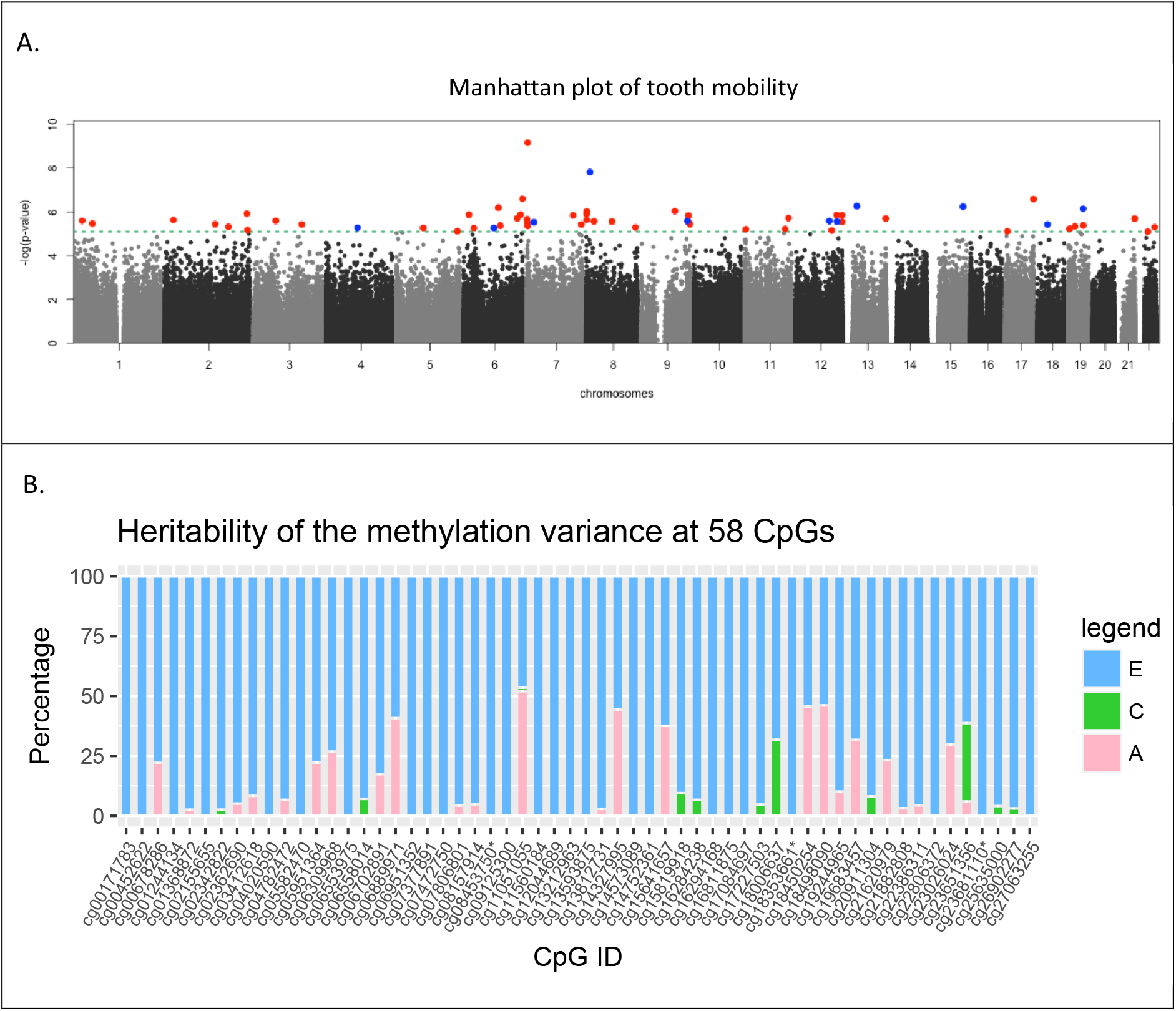
Epigenome-wide association results on tooth mobility. **(A)** Epigenome-wide association results, where green dotted line shows the epigenome-wides significant threshold (8.00e-06). There are 58 points epigenome-wide significant association with tooth mobility, showing positive (red) and negative (blue) correlation with tooth mobility. (B) The percentage of each genetic, shared environmental and unique environmental factor contribution to the variance in methylation levels at the 58 CpG-sites. Pink (A, heritability), green (C, common environment), and blue (E, unique environment) bars represent genetic, shared environmental and unique environmental effects respectively. Three CpGs denoted by * are located in or near to genes that also showed exon expression levels with a nominally significant association with tooth mobility.

Three (cg18353661, cg23681110, cg08453750) out of 58 CpG sites were located in or near to genes that also showed exon expression levels with a nominally significant association with tooth mobility (Figure 2B). Twin-based heritability estimates decomposing the impact of genetic, shared environmental and unique environmental effects on the variance of each blood DNA methylation level showed few strongly heritable sites (heritability > 20% at 12 CpG-sites), while the majority of signals were explained by environmental contributions (Figure 2B). Interestingly, a common environmental component was identified at 12 out of the 58 CpG sites.

We explored if inclusion of technical and biological covariates could attenuate the genomic inflation observed in the EWAS of tooth mobility. When we re-analysed the data using additional participants, that is, including subjects who had dental visits more than 5 years before and after the DNA extraction time (n=717), the genomic inflation factor was attenuated (lambda = 1.14) and only one epigenome-wide significant signal remained. This signal was the top-ranked CpG-site from the set of 58 CpG reported above at cg08157914 in *IQCE* (beta = 0.35, p-value = 1.02-8, FDR = 0.007).

### Enrichment of differential methylation at chronic periodontitis GWAS loci and biological candidates across tissues

We integrated our genome-wide differential methylation profiles in periodontal disease with previously published candidate regions for chronic periodontitis. Altogether, we explored data at 28 regions (256 CpG-sites) that were previously identified either through GWAS efforts or biological studies into periodontal disease (Table 2). We observed evidence for enrichment of epigenetic associations with both periodontal traits in the blood epigenome-wide results (Figure 3A). In blood, 26 CpG sites (tooth mobility) and 15 CpG sites (gingival bleeding) were nominally significantly associated with dental phenotypes. CpG sites that were nominally significantly associated with both periodontal traits mapped to eight genes *(VDR, IL6ST, TMCO6, IL1RN, CD44, IL1B, WHAMM*, and *CXCL1*).

**Figure 3.**
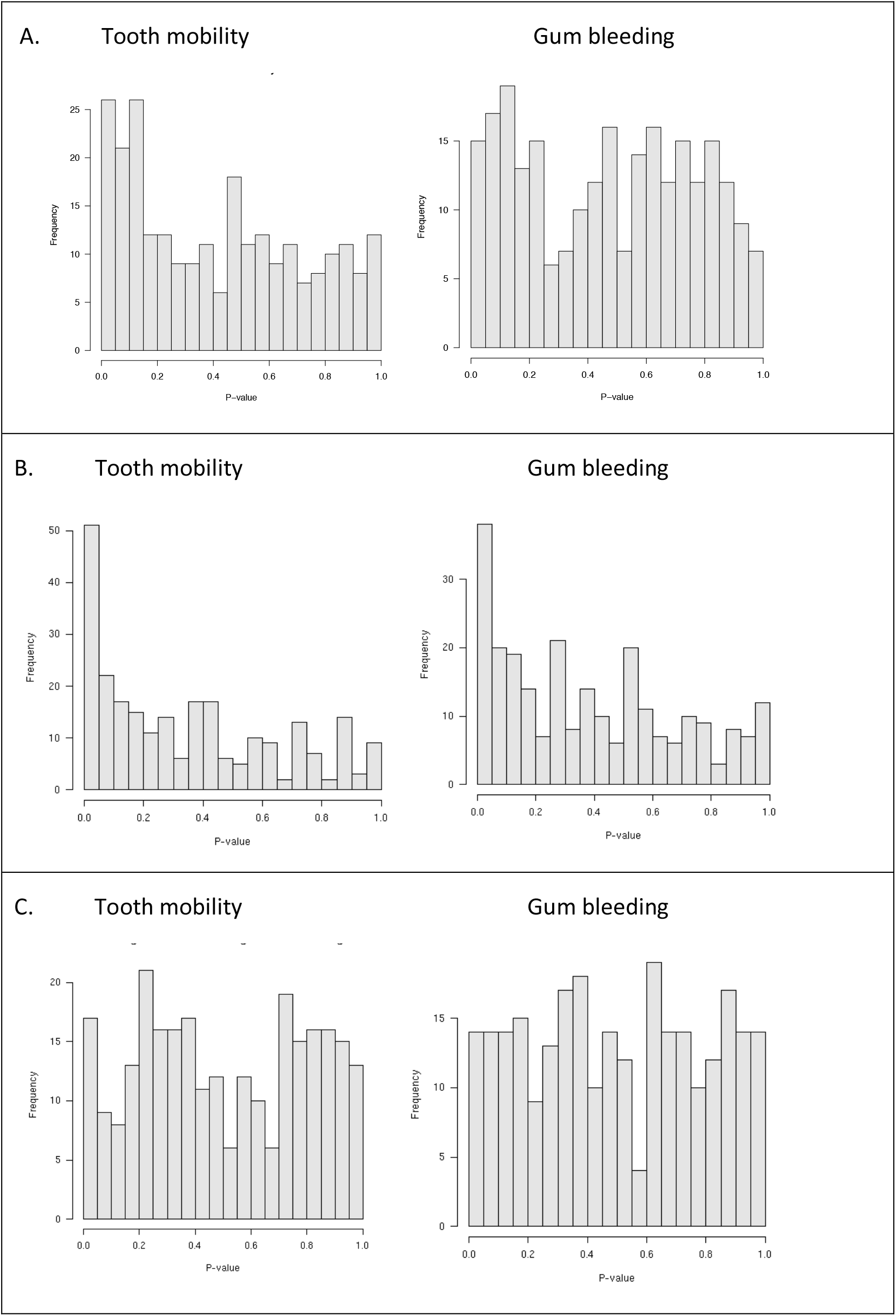
Evidence of methylation-phenotype association at candidate regions for chronic periodontitis across three tissues. The histograms show the strength of association between methylation levels in blood (A), buccal (B) and adipose (C) tissues at CpG-sites mapping to 28 candidate regions and periodontal traits.

To assess the persistence of dental phenotype associations across tissues, we carried out additional epigenetic analyses between both periodontal traits and two further epigenetic datasets from buccal and adipose tissue. In buccal tissue, a stronger phenotype enrichment was observed compared to that in blood (Figure 3B), and 51 (tooth mobility) and 38 (gingival bleeding) CpG sites were nominally associated with the dental phenotypes. CpG sites that were nominally significantly associated with both periodontal traits mapped to 16 genes (*IL1RN, IL1B, CD44, VDR, MMP13, MED24, CCR1, TMCO6, MMP3, IL6ST, TLR4, IL6, IL10, CXCL1, WHAMM* and *SNORD124*), which included all 8 genes detected in blood. In summary, moderate to strong enrichment of dental phenotype association P-values was observed in the blood and buccal epigenetic profiles, while in contrast no enrichment was observed in adipose tissue (Figure 3C).

We further explored the eight genes at which nominally significant differential methylation was detected in both blood and buccal tissue across both tooth mobility and gingival bleeding phenotypes. First, we carried out tests of association between the phenotype-associated blood DNA methylation signals and gene expression levels at these eight genes within the linear mixed effects regression framework. We found Bonferroni significant associations (P = 0.00625) between periodontal traits and exon expression levels for exons in *WHAMM* (exon ENSG00000156232.5_83499351_83499831: beta (SE) = 2.42e-6 (7.83e-7); p-value = 2.2e-3, exon ENSG00000156232.5_83495140_83495248: beta (SE) = 2.68e-6 (9.13e-7); p-value = 3.6e-3, exon ENSG00000156232.5_83477973_83479087: beta (SE) = 2.73e-6 (9.36e-7); p-value = 3.9e-3) and *TMCO6* (exon ENSG00000113119.8_140023365_140023551: beta (SE) = −0.37 (0.13); p-value = 6.0e-3). Second, to further explore their functional contribution, we examined the association between the methylation levels of the phenotype-associated CpG-sites in these two genes (cg00115297 in *WHAMM* and cg00916199 in *TMCO6*) and fasting blood metabolites profiles. Nominally significant associations with methylation levels at these CpGs were observed for 48 metabolites (Figure 4), with the most-associated signals for lipid related metabolites. Six out of 23 identified metabolites in a nominally significant association with *WHAMM* methylation were related to phosphocholine metabolism and the other three were sulfate metabolites. With respect to the association with *TMCO6* methylation, 8 out of 25 identified metabolites were related to carnitine.

**Figure 4.**
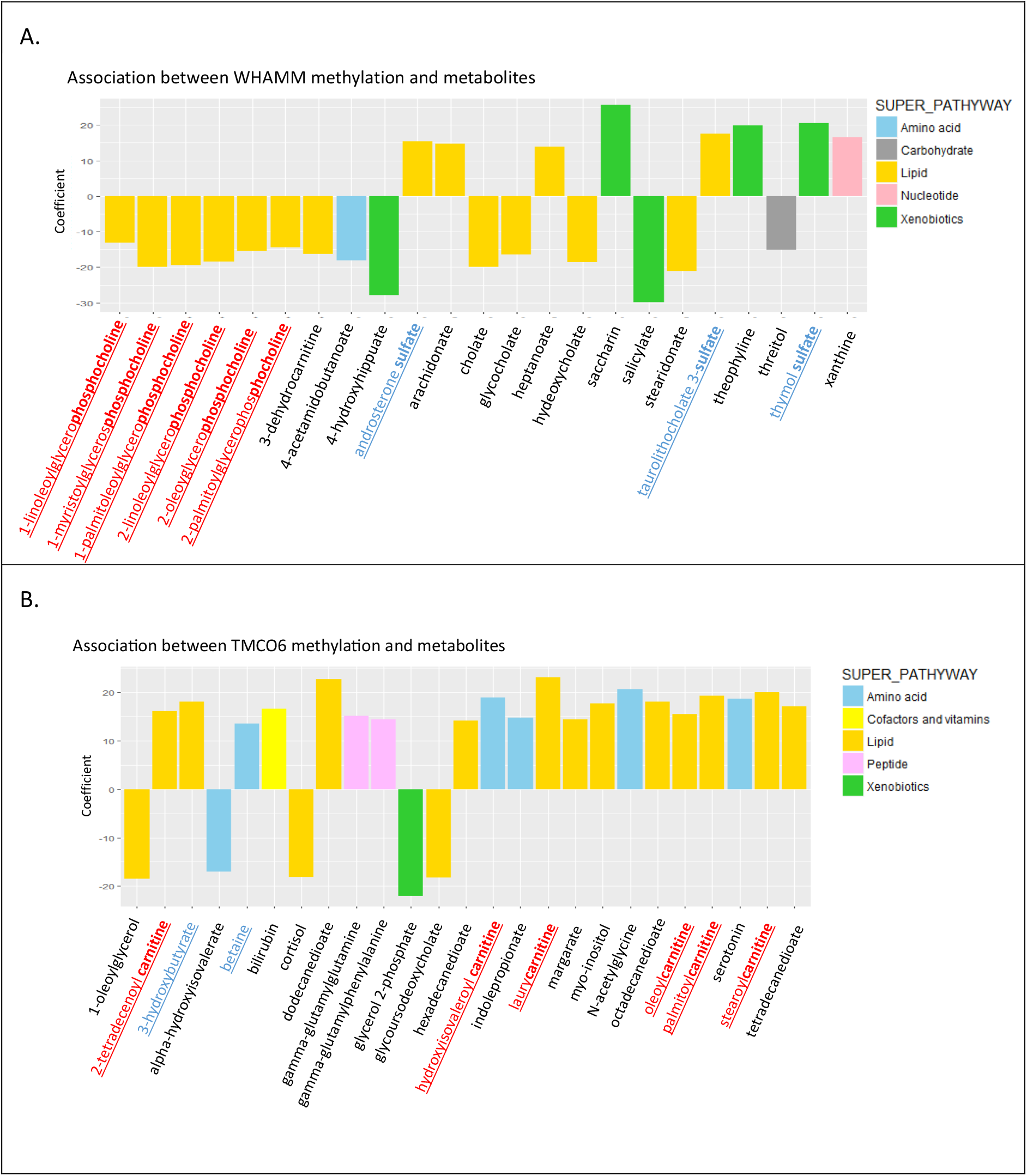
Functional metabolomic characterization of *WHAMM* and *TMCO6* methylation signals. Blood metabolites showed a nominally significant association with blood methylation levels at cg00115297 proximal to *WHAMM* **(A)** and at cg00916199 proximal to *TMCO6* **(B)**. The y axis shows the coefficient of the association between methylation and metabolites level.

## DISCUSSION

We present the first large-scale epigenome-wide study of periodontitis. Our results identified differential methylation at specific CpG sites between periodontal trait positive and negative individuals, which suggests that epigenetic changes may help explain the variation in the susceptibility to chronic periodontitis. Our findings detected hypo-methylation in the 5’UTR of *ZNF804A* and corresponding up-regulation of gene expression in individuals who experience gingival bleeding. These data suggest that *ZNF804A* should be considered as a candidate gene related to genomic regulation and gingival inflammation. Blood metabolomics profile analysis highlighted eight metabolites, including ornithine and carnitine, which were significantly associated with the differential methylation signal in *ZNF804A*. Ornithine was reported to play an important role on metabolic cross-feeding between *Streptococcus gordonii* and *Fusobacterium nucleatum* [25]. *S. godonii* is considered to express an arginine-ornithine antiporter, which potentially could influence the inter-bacterial communication with periodontal pathogens. Kuboniwa’s group also reported that the level of ornithine in saliva showed a decreasing trend after an intervention of debridement [26]. Taken together, these findings suggest a potential sequence of events in early stage periodontal disease that may lead towards disease progression, where in early stage periodontal disease - where gingival inflammation is observed - hypomethylation of cg21245277 may up-regulate the expression of *ZNF804A*, which in turn could elevate the level of ornithine in blood.

In terms of more severe stages of periodontal disease, where tooth mobility or self-reported history of gum disease is a more appropriate phenotype marker, we identified 58 CpG sites with epigenome-wide significant signals, although the observed genomic inflation may result in an increase in the number of false positive findings that we report here [27]. The most strongly associated signal for this phenotype was in the gene body of the *IQCE* gene and was hyper-methylated in individuals who presented with tooth mobility, the severe stage of periodontitis. However, there was not a significant signal for differential gene expression in this gene. Gene expression patterns associated with methylation levels at tooth-mobility associated signals were obtained at 3 CpG sites (out of 58 CpG-sites) and in three genes (*MAD1L1, TRAPPC9*, and *OSBPL5*). Epigenetic regulation may impact gene expression levels at these genes in severe cases of periodontal disease. According to the heritability findings for the variance of blood DNA methylation levels, 21% of the 58 CpG sites showed more than 20% genetic influence, and similarly 23% of the variance of the methylation level on the CpG site related to *ZNF804A* gene was explained by a genetic component. This finding is consistent with previous reports that DNA methylation levels can be under genetic control [28].

The most striking finding of our results was generated from a candidate gene approach. We identified eight genes (*VDR, IL6ST, TMCO6, IL1RN, CD44, IL1B, WHAMM*, and *CXCL1*) that displayed methylation associations with both periodontal traits across blood and buccal tissues. Of these, *WHAMM* and *TMCO6* were identified as likely signals of genomic regulation in periodontal disease, due to significant association between phenotype-associated methylation and gene expression levels at these genes. Both findings suggest that *WHAMM* and *TMCO6* are strong candidates to shed light on epigenetic modifications over the course of periodontal disease. *WHAMM* encodes a protein which regulates the membrane dynamics and functions at the interface of microtubule and actin cytoskeletons. This gene was previously identified as a candidate from two different GWAS studies in chronic periodontitis [1, 29]. *TMCO6* is located near *CD14*, which is an important contributor to periodontal inflammation as it encodes a protein which is part of the innate immune system and also recognizes lipopolysaccharides (LPS) components of gram negative bacterial cell wall. Several case-control studies also found increased serum concentration of *CD14* among the patients with periodontitis [30, 31].

Further analysis with blood metabolites showed that the epigenetic modifications at both *WHAMM* and *TMCO6* were related to lipid metabolite biological pathways. Lipid metabolism has been well studied in microbial organisms including in periodontal pathogenic bacteria such as *P. gingivalis* and *A. actinomycemcomitans* [32] [33] [34]. Approximately 30% of the metabolites, associated with *TMCO6* methylation were related to carnitine metabolism, which can be utilized by gram (+) (-) bacteria [35]. Three out of 23 identified metabolites related to *WHAMM* methylation levels were sulfates. Sulfate is known as a crucial biological markers to evaluate periodontal disease as it is one of the chemical components of glycosaminoglycans [36], which are generally elevated under inflammatory condition.

Our study has a number of limitations. Firstly, we used self-reported data to assess periodontal disease status. This is likely to underestimate the presence of disease compared to the observational data including periodontal pocket depth, attachment loss and bleeding on probing in person by dental professionals. Our previous validation study of the self-reported data, which included 80 individuals, showed that self-reported history of gum disease or tooth mobility had low sensitivity, 0.28, but a very high specificity and positive predictive value of 1. This result suggests that there is a high degree of certainly about the periodontitis phenotype in test subjects when compared against a control group of unknown periodontal phenotype, which arguably decreases the power of the study to detect a true difference between groups, but makes a type I error unlikely. This is consistent with previous studies of self-reported periodontitis where questions assessing loose teeth and professional diagnosis of gum disease are good predictors of periodontitis compared to other questions, but again, as with our study, with low sensitivity [37, 38]. For gingival bleeding there was sensitivity of 0.48 and specificity of 0.75 respectively. There is very little existing data that demonstrate the reliability of self-reported gingival bleeding and thus our results with the gingival bleeding trait should be treated with caution, particularly when considering the relatively poor specificity seen in our own validation.

Another limitation of our periodontal information is the time difference between DNA extraction time and dental phenotype collection, and dental questionnaires which capture the history of periodontal conditions rather than current status. However, we excluded participants for whom the time difference between DNA extraction and dental questionnaires was over 5 years. We reasoned that subjects are likely to suffer from the same periodontal condition for roughly 5 years, because periodontal disease encompasses the nature of chronic inflammation.

Another limitation is our ability to address tissue specificity in disease relevant tissues. Epigenetic changes are tissue specific and patterns in blood may not be shared across tissues. Tissues within oral cavity such as gingiva, buccal mucosa and saliva are considered preferable to examine the regulatory genomic effects in periodontal disease due to their involvement or proximity to disease-affected areas [39]. We attempted to address this point by exploring methylation profiles in a modest sample of buccal tissue.

Although all of these drawbacks need to be further considered, our notable findings from the epigenome-wide analysis and candidate gene approach identify novel genomic regions in periodontitis and highlight candidate genes for further research. Previous studies examining epigenetic variation related to periodontitis have applied the case control design with moderate samples and limited information on specific factors that may influence on DNA methylation such as age, gender, smoking status and technical covariates. Overall, the current study suggests that epigenetic modifications may play an important role in the susceptibility and progression of periodontal disease. Future studies could focus on identifying environmental factors, which trigger DNA methylation changes relevant to periodontitis. Periodontal pathogenic bacteria have the potential to cause alteration in cellular DNA methylation status. Moreover, environmental factors such as aging, smoking and stress can have impacts on epigenetic modifications. Further future work could explore in more depth the tissue specificity of epigenetic modifications in periodontitis. Tissue-shared epigenetic modifications are valuable as blood methylation data are available in other cohorts, which allows us to conduct replication studies.

## CONCLUSIONS

Epigenome-wide analyses and a candidate gene approach in adult female twins identified differentially methylated signals in periodontal disease. In conjunction with transcriptomics and metabolomics analyses, we conclude that the epigenetic changes may have functional regulatory effects over the course of development and progression of periodontal disease.

## LIST OF ABBREVIATIONS

EWAS: Epigenome-wide association scans
GWAS: Genome-wide association scans
FDR: False Discovery Rate
SNPs: Single nucleotide polymorphism
MZ: Monozygotic
DZ: Dizygotic
BMIQ: Beta mixture quantile normalization
DMPs: Differentially methylated positions
LMER: Linear mixed effects regression
LPS: Lipopolysaccharide

## DECLARATIONS

### Ethics approval and consent to participate

Ethical approval was granted by the National Research Ethics Service London-Westminster, the St Thomas’ Hospital Research Ethics Committee (EC04/015 and 07/H0802/84). All research participants have signed informed consent prior to taking part in any research activities.

### Consent for publication

All authors have read and approved the manuscript for publication.

### Availability of data and material

The majority of the datasets analysed in the current study are available through GEO GSE62992 (blood methylation), ArrayExpress E-MTAB-1866 (adipose methylation), and EGA EGAD00001001088 (blood expression). Additional individual-level data are not permitted to be shared or deposited due to the original consent given at the time of data collection. However, access to these metabolite and phenotype data can be applied for through the TwinsUK data access committee. For information on access and how to apply http://www.twinsuk.ac.uk/data-access/submission-procedure/.

### Competing interests

The authors declare no conflict of interest.

### Funding

The study received support from the ESRC (ES/N000404/1 to J.T.B) and the JSPS (the Postdoctoral Fellowship for Research Abroad to Y.K.). The TwinsUK study was funded by the Wellcome Trust; European Community’s Seventh Framework Programme (FP7/2007–2013); National Institute for Health Research (NIHR)- funded BioResource, Clinical Research Facility and Biomedical Research Centre based at Guy’s and St Thomas’ NHS Foundation Trust in partnership with King’s College London. SNP genotyping was performed by The Wellcome Trust Sanger Institute and National Eye Institute via NIH/CIDR.

## Acknowledgments

The authors would like to thank Dr Tiphaine Martin for assistance with CoMET.

## Authors’ contributions

Y.K. and J.T.B. designed the study and outlined the main conceptual ideas. J.T.B. and C.S. generated the primary datasets, and M.I. and F.H. helped guide data generation and results interpretation. J.T.B., K.S.S., and C.S. oversaw data analysis. Y.K. lead data analysis. P.C-T, J.E.C-F. A.D.S.C.A., J.S.E-S.M., C.L.R., and A.V. contributed data analysis. J.T.B. and Y.K. wrote the article and all authors provided critical feedback and helped shape the research, analysis and manuscript.

